# MethyLasso: a segmentation approach to analyze DNA methylation patterns and identify differentially methylation regions from whole-genome datasets

**DOI:** 10.1101/2023.07.27.550791

**Authors:** Delphine Balaramane, Yannick G. Spill, Michaël Weber, Anaïs Flore Bardet

## Abstract

DNA methylation is an epigenetic mark involved in the regulation of gene expression and patterns of DNA methylation anticorrelates with chromatin accessibility and transcription factor binding. DNA methylation can be profiled at the single cytosine resolution in the whole genome and has been performed in many cell types and conditions. Computational approaches are then essential to study DNA methylation patterns in a single condition or capture dynamic changes of DNA methylation levels across conditions. Towards this goal, we developed MethyLasso, a new approach based on the segmentation of DNA methylation data, that enables the identification of low-methylated regions (LMRs), unmethylated regions (UMRs), DNA methylation valleys (DMVs) and partially methylated domains (PMDs) in a single condition as well as differentially methylated regions (DMRs) between two conditions. We performed a rigorous benchmarking comparing existing approaches by evaluating the number, size, level of DNA methylation, boundaries, CpG content and coverage of the regions using several real datasets as well as the sensitivity and precision of the approaches using simulated data and show that MethyLasso performs best overall. MethyLasso is freely available at https://github.com/abardet/methylasso.

## INTRODUCTION

DNA methylation is an epigenetic mark that consists in the addition of a methyl group to cytosines mainly in the context of cytosine-guanine (CpG) dinucleotides. It is involved in the regulation of gene expression as it is able to block transcription factors from binding to DNA (1, 2). Patterns of DNA methylation anti-correlate with transcription factor binding where most of the inactive genome is fully methylated and active regulatory regions bound by transcription factors are lowly-or unmethylated (LMRs or UMRs) (3, 4). Large unmethylated regions that often mark developmental genes were termed DNA methylation valleys (DMVs) (5, 6) and partially methylated domains (PMDs) have been described as large domains with heterogeneous methylation levels in immortalized cell lines and cancer samples (7, 8). DNA methylation is a reversible mark and demethylation was shown to be induced by transcription factor binding (3, 9, 10). DNA methylation patterns are therefore dynamic notably during cellular differentiation throughout development and disease conditions such as cancer.

Several experimental approaches have been developed to profile DNA methylation patterns (11). Bisulfite conversion (12) and more recently enzymatic conversion (13) followed by high-throughput sequencing can be applied to profile whole-genome DNA methylation at single cytosine resolution (Bis-seq or EM-seq respectively). Computational approaches are then essential to study DNA methylation patterns in a single condition or capture dynamic changes of DNA methylation levels across conditions.

The identification of hypomethylated regions in a single condition is of great interest as it can be used to predict active regulatory regions bound by transcription factors. MethylSeekR (4) is the most widely used tool to identify LMRs, UMRs and PMDs. In order to circumvent the uncertainty of methylation levels due to the low sequencing coverage of individual CpGs, MethylSeekR smooths methylation levels over three consecutive CpGs with a minimal coverage of 5 reads. It then identifies hypomethylated regions as stretches of consecutive CpGs with methylation below 50%. Since it expects LMRs to be located at CpG-poor regions and UMRs to be found at CpG-rich regions called CpG islands, MethylSeekR uses CpG content as a threshold to further classify regions as LMRs (e.g. below 30 CpGs) or UMRs (e.g. above 30 CpGs). False discovery rate (FDR) is computed using shuffled CpG methylation levels. PMDs are identified using a Hidden Markov model with sliding windows of 100 consecutive CpGs and classified according to the methylation levels.

The identification of differentially methylated regions (DMRs) to compare conditions is one of the main analyses performed when investigating DNA methylation patterns. Many tools with various approaches have been developed for whole-genome methylation datasets that have been reviewed and benchmarked by several studies (14–17). Most approaches first rely on the identification of single differentially methylated CpGs (DMCs) and then aggregates them into DMRs. One of the first approach developed to identify DMRs is BSmooth (18), which applies local likelihood smoothing to overcome potential biases due to low coverage. It requires replicates to calls DMCs using a t-test and group the ones above a specific threshold into DMRs. RADmeth (19) is based on a beta-binomial regression approach. It calls significant DMCs using a log-likelihood ratio test and neighboring DMCs are combined using a weighted Z-test to obtain DMRs. DSS (20) is based on a Bayesian hierarchical model based on the beta-binomial distribution. It can also smooth the methylation values, calls significant DMCs using a Wald test and groups them as DMRs using thresholds such as p-value, minimum length and minimum number of CpGs. Defiant (21) employs an approach where the sample’s variance is weighted based on coverage. It calls significant DMCs using a Welch’s t-test if several replicates are available or a Fisher’s exact test otherwise and uses a weighted Welch expansion to identify DMRs. Dmrseq (22), that now replaces BSmooth, also performs smoothing and uses the same beta-binomial approach than DSS to call significant DMCs. It then uses a continuous autoregressive correlation to identify DMRs and controls for FDR using the Benjamini and Hochberg procedure. DMRcate (23) that was first developed for methylation array data was recently adapted for whole genome Bis-seq data and uses the limma to generate per-CpG t-statistics and kernel smoothing and groups them into DMRs using a FDR threshold.

Only few approaches define DMRs directly based on the changes of DNA methylation differences without any assumption about the data distribution. Metilene (24) performs a first pre-segmentation of the genome and then a circular binary segmentation approach is used to iteratively reduce the size of the regions to maximize the mean DNA methylation difference. Segmentation stops when a minimum of CpGs is reached or when the two-dimensional Kolmogorov–Smirnov p-value does not improve. Adjusted p-values are also provided as well as p-values from a Mann–Whitney U test.

Although many tools have been developed to identify DMRs, the regions identified differ in terms of number, level of DNA methylation differences and region boundaries and there is limited overlap between DMRs from different approaches.

Here, we present MethyLasso, a new segmentation approach to analyze DNA methylation patterns and identify differentially methylation regions from whole-genome datasets. MethyLasso models DNA methylation data using a regression framework known as generalized additive model. It relies on the fused lasso to estimate regions in which the methylation is constant. We apply MethyLasso on data from a single condition to identify LMRs, UMRs and DMVs based on DNA methylation levels independently of CpG content as well as PMDs. We also adapted MethyLasso to identify DMRs between two conditions. We compare MethyLasso to other methods that were shown to perform best in the literature and show that MethyLasso outcompetes them.

## MATERIAL AND METHODS

### Input data format

MethyLasso runs on whole genome DNA methylation data (e.g. Bis-seq or EM-seq) where DNA methylation is measured for each cytosine in the genome, which can then be summed for each CpG. It should not be applied to reduced representation bisulfite sequencing (RRBS) data because the segmentation would group distant cytosines separated by cytosines with no coverage. The data takes the form of a matrix containing for each C or CpG position in the genome (chromosome, start, end), the percent of methylated sequences (meth) out of the total number of sequences covering this position (cov). This can alternatively be calculated from the number of methylated Cs (mC) and the number of unmethylated Cs (uC). By default, data in a Bismark output format (25) is used (chromosome, start, end, percent_methylation, count_methylated, count_unmethylated but other formats can also be specified (see MethyLasso README). MethyLasso only keeps by default positions covered by at least 5 reads (program argument -c).

### MethyLasso general framework

MethyLasso models DNA methylation data using a nonparametric regression framework known as a Generalized Additive Model. It relies on the fused lasso method (26) to segment the genome by estimating regions in which methylation is constant. This model is adapted from the one applied for Hi-C data in Binless normalization (27). For a single whole genome DNA methylation dataset, the resulting data takes the form of a N-dimensional vector of the observed methylated fraction y=M/C from methylated counts M (program argument -mC), and total sequencing coverage C (program argument --cov). y is given a normal likelihood with unknown true methylation fraction β. A 1D fused lasso prior is then placed on β, resulting in a minus log posterior proportional to the L1-penalized weighted least squares target. *λ*_2_ corresponds to the strength of the fusion.

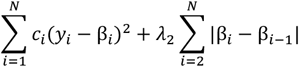

We model one set of coefficients β by condition. In the case of multiple replicates per condition, or multiple conditions, a design matrix X which maps replicates to conditions is built and the mean of y is then Xβ. We model each chromosome independently. The resulting maximum posterior estimate β is a piecewise constant function, which is then used as a basis to construct a segmentation of the methylation data.

### Identification of low methylated regions (LMRs), unmethylated regions (UMRs) and DNA methylation valleys (DMVs)

#### Segmentation of the genome

For each single condition (possibly with multiple replicates), MethyLasso performs a fused lasso segmentation of the methylation values with λ_2_ set to 25 to be able to identify small regions of constant methylation. We obtain a vector of methylated fractions β, with one coefficient per CpG. This vector is piecewise constant and forms a first crude segmentation of the methylome. Segments are split when they contain gaps in between two CpGs of 500 bp or more. Their borders are adjusted to start and end on a CpG. Then, for each segment, we compute the number of CpGs, mean methylation and standard deviation.

#### Identification of LMRs and UMRs

We first define admissible segments if they have a minimum size of 10 bp and a minimum of 4 CpGs (program argument -n). Among admissible segments, we define segments of interest as those whose methylation is smaller than that of its nearest admissible neighbors (on both sides). We then define flanking segments as admissible segments with a minimum size of 300 bp, which are immediate neighbors to a segment of interest but are not segments of interest themselves. Candidate LMRs are defined as segments of interest which are separated from their flanking segments by at least 2 standard deviations, whose mean methylation is below 0.5 and whose flanking segments are no further away than 5,000 bp. Candidate LMRs that simultaneously have a CpG count below 10 and flanking segments with a methylation below 0.5 are not retained in the final LMR call. Candidate LMRs which have a mean methylation below 0.1 and simultaneously whose standard deviation is smaller than 0.1 are instead called UMRs. Remaining candidate LMRs constitute the final LMR call. Finally, among segments of interest with a mean methylation below 0.1 and standard deviation below 0.1 which were not part of the candidate LMR set, we declare those which have a size above 500 bp and more than 30 CpGs/kb as additional UMRs. LMRs or UMRs which overlap a DMV (see below) and LMRs which overlap a PMD (see below) are not reported.

#### Identification of DMVs

We further group segments whose mean methylation is continuously below 0.1 as long as they are not separated by a large gap of 5 kb. Among those merged segments and already defined UMRs, we define DMVs if their size is equal or greater than 5 kb. We then recompute their number of CpGs, mean methylation and standard deviation. We exclude DMVs that have an average distance between CpGs larger than 500 bp.

#### Output files

MethyLasso outputs one file per condition containing the genomic coordinates and characteristics of the regions (chromosome, start, end, number of CpGs, mean methylation, standard deviation and annotation as LMRs and UMRs including DMVs). It also generates plots showing histograms of the reads sequencing depth and CpG methylation from positions with at least 5 reads (see MethyLasso README).

### Identification of partially methylated domains (PMDs)

#### Segmentation of the genome

For each single condition (possibly with multiple replicates), MethyLasso performs a fused lasso segmentation of the standard deviation with λ_2_ set to 1000 to identify large regions where methylation varies widely. Segments are split when they contain gaps in between two CpGs of 5 kb or more or when they contain a UMR or DMV. Then, for each segment, we compute the number of CpGs, mean methylation and standard deviation.

#### Identification of PMDs

We first define admissible segments if they have a size larger than 5 kb, a minimum of 4 CpGs and an average distance between CpGs smaller than 500 bp. PMDs are then defined as admissible segments with a mean methylation below 0.7 and a standard deviation above 0.15. Contiguous PMDs are then merged into a single region. We then recompute their number of CpGs, mean methylation and standard deviation.

#### Output files

MethyLasso outputs one file per condition containing the genomic coordinates and characteristics of the regions (chromosome, start, end, number of CpGs, mean methylation, standard deviation and annotation as PMDs).

### Identification of differentially methylated regions (DMRs)

#### Segmentation of the genome

In presence of two conditions (possibly with multiple replicates), the statistical model is designed such that one set of coefficients represents the methylation of the reference condition, while the remaining sets of coefficients represent the methylation difference of the remaining condition to the reference. Each set of coefficients is obtained by a 1D fused lasso as described with λ_2_ set to 25 to be able to identify small regions, and yields a segmentation of the methylation differences between a condition and the reference. Then, for each segment, we compute the number of CpGs, mean methylation difference and standard deviation.

#### Identification of DMRs

We group neighboring segments when their mean methylation difference is below 0.1 (program argument -d). Their borders are adjusted so that they start and end on a CpG. We then recompute their number of CpGs, mean methylation difference and standard deviation. We also compute a coverage score for each DMR as the percent of CpGs covered in all samples (conditions and replicates) and discard DMRs with a coverage score below 70%. Finally, we calculate the Wilcoxon p-value for each DMR by comparing the methylation difference at each CpG in the two conditions. False discovery rate (FDR) is controlled using the Benjamini-Hochberg procedure on the p-values. We retain DMRs with a minimum of 4 CpGs (program argument -n), a methylation difference above or equal to 0.1 (program argument -d), a p-value below or equal to 0.05 (program argument -p). Alternatively, DMRs can also be selected according to the FDR threshold (program argument -q).

#### Annotation of DMRs using LMRs, UMRs, DMVs and PMDs

If LMRs, UMRs, DMVs and PMDs were defined in each condition, DMRs will be annotated according to those annotations. DMRs are annotated in each condition by the regions with which it overlaps most if any and at least 10% for LMRs, UMRs and DMVs and 50% for PMDs. Change of annotation is then indicated as from the reference condition to the other e.g. LMRtoUMR, (nothing)toPMD, fromDMV(to nothing).

#### Output files

MethyLasso outputs one file containing the genomic coordinates and characteristics of the regions (chromosome, start, end, number of CpGs in each condition, coverage score, mean methylation in each condition, mean methylation difference, p-value, FDR and annotation). It also generates a scatterplot showing the methylation level of the DMRs in the reference condition compared to the other condition.

### MethyLasso benchmarking

#### DNA methylation datasets

We used whole genome Bis-seq datasets publicly available in the NCBI gene expression omnibus (GEO) repository from human embryonic stem cells (ESH1) and fetal lung fibroblasts cells (IMR90) (28) (GEO accessions GSM429321, GSM429322, GSM429323 GSM432685 and GSM432686 for ESH1 one replicate, GSM432687, GSM432688 and GSM432689 for IMR90 replicate one, and GSM432690, GSM432691 and GSM432692 for IMR90 replicate two), primary human colorectal cancer and normal cells (29) (GEO accessions GSM1204465 and GSM1204466 respectively with three replicates each), and mouse hematopoietic stem cells (HSC) and multipotent progenitor populations (MPP) (30) (GEO accessions GSM1274424, GSM1274425 and GSM1274426 for HSC three replicates and GSM1274433, GSM1274434 and GSM1274435 for MPP three replicates).

#### Data processing

Raw sequencing reads were trimmed using trim_galore (version 0.6.4 options -q 20 --stringency 2) (http://www.bioinformatics.babraham.ac.uk/projects/trim_galore/) and mapped to the human genome reference hg38 or the mouse mm10 using bismark (version 0.22.1) (25). Non-converted and duplicated reads were filtered out using filter_non_conversion –percentage_cutoff 50 –minimum_count 5 and deduplicate_bismark. Methylation levels were extracted using bismark_methylation_extractor. Methylation was summarized per CpG by overlapping the methylation levels per cytosine with CpGs of the reference genome using a custom script.

#### Simulated DMRs

We used whole genome Bis-seq data containing simulated DMRs from Metilene to evaluate the performance of MethyLasso and other methods. They simulated DMRs with two different backgrounds (homogeneous background 1 and heterogeneous background 2), each with four subsets, from the largest methylation difference to the smallest. Each subset has ten replicates. Data is available from http://www.bioinf.uni-leipzig.de/Software/metilene/Downloads/.

#### Comparison with other methods

For the identification of LMRs, UMRs and PMDs, we compared MethyLasso to MethylSeekR (version 3.6.1) (4). MethylSeekR was executed with default settings. Since MethylSeekR does not identify DMVs, we called our DMVs as UMRs when comparing UMRs. In order to compare methylation levels in LMRs, UMRs and PMDs from the two different programs, we recomputed them by first summing the counts from the replicates for each condition to calculate the methylation level per CpG and then calculating the mean methylation in the region.

For the identification of DMRs, we compared MethyLasso to six other programs: Defiant (1.1.9) (21), Dmrseq (1.6.0) (22), DSS (2.34.0) (20), RadMeth (19), DMRcate (2.14.0) (23) and Metilene (0.2-8) (24). We used those programs with default setting except for a minimum coverage of 5 reads per CpG, a minimum of 4 CpGs and at least 10% methylation change to call DMRs. DSS was run with smoothing since they always recommend to smooth data for whole-genome Bis-seq data. We also ran Dmrseq, RadMeth, Metilene and DMRcate with more relaxed thresholds in order to obtain the best 150,000 DMRs for colorectal cancer vs. normal cells, best 270,000 DMRs for ESH1 vs. IMR90 data and best 5,000 for HSC vs. MPP. In order to compare methylation differences in DMRs from the different programs, we recomputed them by first summing the counts from the replicates for each condition to calculate the methylation level per CpG, then calculating the difference between two conditions and finally calculating the mean difference in the region. The DMRs identified by the different methods are available to load on the UCSC Genome Browser (My Data / Track Hubs) using the following URL https://g-948214.d2cf88.03c0.data.globus.org/hub_methylasso.txt.

#### Computational setup

Benchmarking was performed on a server using an Intel Xeon Silver CPU (10 cores, 20 threads, 2.2 GHz and 96GB RAM) and MethyLasso can also run on a MacBook Air with a 1,6 GHz Intel Core i5 processor and 8 GB of memory.

#### Data analyses

All genomic data analyses were performed using bash scripts using the command awk and bedtools (31) and R for plots and statistics. The upset plots we generated using the intervene R package (32).

### Software implementation and availability

MethyLasso’s segmentation code is written in C++ and the regions’ identification code in R. MethyLasso’s code and manual as well as the code developed to benchmark the methods are available on github at https://github.com/abardet/methylasso.

## RESULTS

### Identification of low-methylated regions (LMRs), unmethylated regions (UMRs) and DNA methylation valleys (DMVs)

MethyLasso is a method that first analyzes DNA methylation patterns in each experimental condition independently. It relies on a fused lasso approach to segment the genome by estimating regions in which the methylation is constant (**see Methods**). It fits the model to DNA methylation data, segments the data and calls different regions such as LMRs (10 to 50% methylation), UMRs (0 to 10% methylation) or DMVs (UMRs larger than 5 kb) according to specific thresholds (**Figure 1A**) (**see Methods**). One advantage over MethylSeekR is that MethyLasso can integrate replicates when available to call each set of regions only once per condition. On data from human IMR90 fibroblast cells (28), MethyLasso identified 29,914 LMRs, 28,308 UMRs including 240 DMVs where MethylSeekR identified more LMRs but less UMRs in both replicates independently (**Figure 1B and Supplementary Figure 1** for other datasets). As expected, UMRs often located at CpG island promoters are larger than LMRs (**Figure 1C**). We observe that LMRs are more often identified by only one of the approaches than UMRs (**Figure 1B**), which can be explained by the fact that UMRs have more CpGs with DNA methylation levels constrained within 0 to 10% whereas LMRs can have a broader middle range of methylation levels (10 to 50%) with higher variance so are expected to be more difficult to identify. When looking at the number of CpGs versus the mean methylation level in each of the regions (UMRs + LMRs), we observe that they cluster as expected: regions with high CpG density such as CpG islands are mostly unmethylated and regions with low CpG density have higher methylation levels (**Figure 1D,E and Supplementary Figure 1**). However, we decided not to include the CpG density as a main parameter to identify UMRs and LMRs as some regions with few CpGs can be completely unmethylated (**Figure 1D**, bottom left corner). Therefore, MethyLasso only sets the threshold between LMRs and UMRs according to their DNA methylation level matching the original definition (3) from 0 to 10% for UMRs and from 10 to 50% for LMRs independently of their CpG density (**Figure 1D** black line). In contrast, MethySeekR sets the threshold between LMRs and UMRs according to their CpG density (30 CpGs for IMR90 data) (**Figure 1E** black line) and therefore some UMRs predicted by MethylSeekR are not completely unmethylated whereas some LMRs are (**Figure 1E**). Since MethylSeekR UMRs correspond to regions dense in CpGs, it also explains why they are larger than MethyLasso UMRs (**Figure 1C**). Therefore, because of the difference of definition of the regions, 8,913 MethyLasso UMRs are identified as LMRs by MethylSeekR and 1,481 LMRs as UMRs (**Figure 1B,F and Supplementary Figure 1**). Even when both programs agree on calling UMRs or LMRs, their region boundaries still differ and even depend on the replicate for MethylSeekR (**Figure 1G**). To evaluate this systematically, we compared the methylation of CpGs within UMRs or LMRs to the CpGs immediately upstream or downstream (**Figure 1H,I**). For UMRs, we observe that both MethyLasso and MethylSeekR show significant shifts of DNA methylation levels between neighboring CpGs across boundaries. However, MethyLasso identifies smaller regions (**Figure 1C**) that are completely unmethylated whereas MethylSeekR includes surrounding CpGs that have higher methylation levels without being fully unmethylated (**Figure 1H**) that might represent CpG island shores (33). For LMRs, we observe that CpGs across boundaries have better shifts in DNA methylation level for MethyLasso than for MethylSeekR and that CpGs within MethyLasso LMRs have more homogenous DNA methylation levels (**Figure 1I**). Similar findings were obtained in several other human or mouse Bis-seq datasets (**Supplementary Figure 2**).

**Figure 1.**
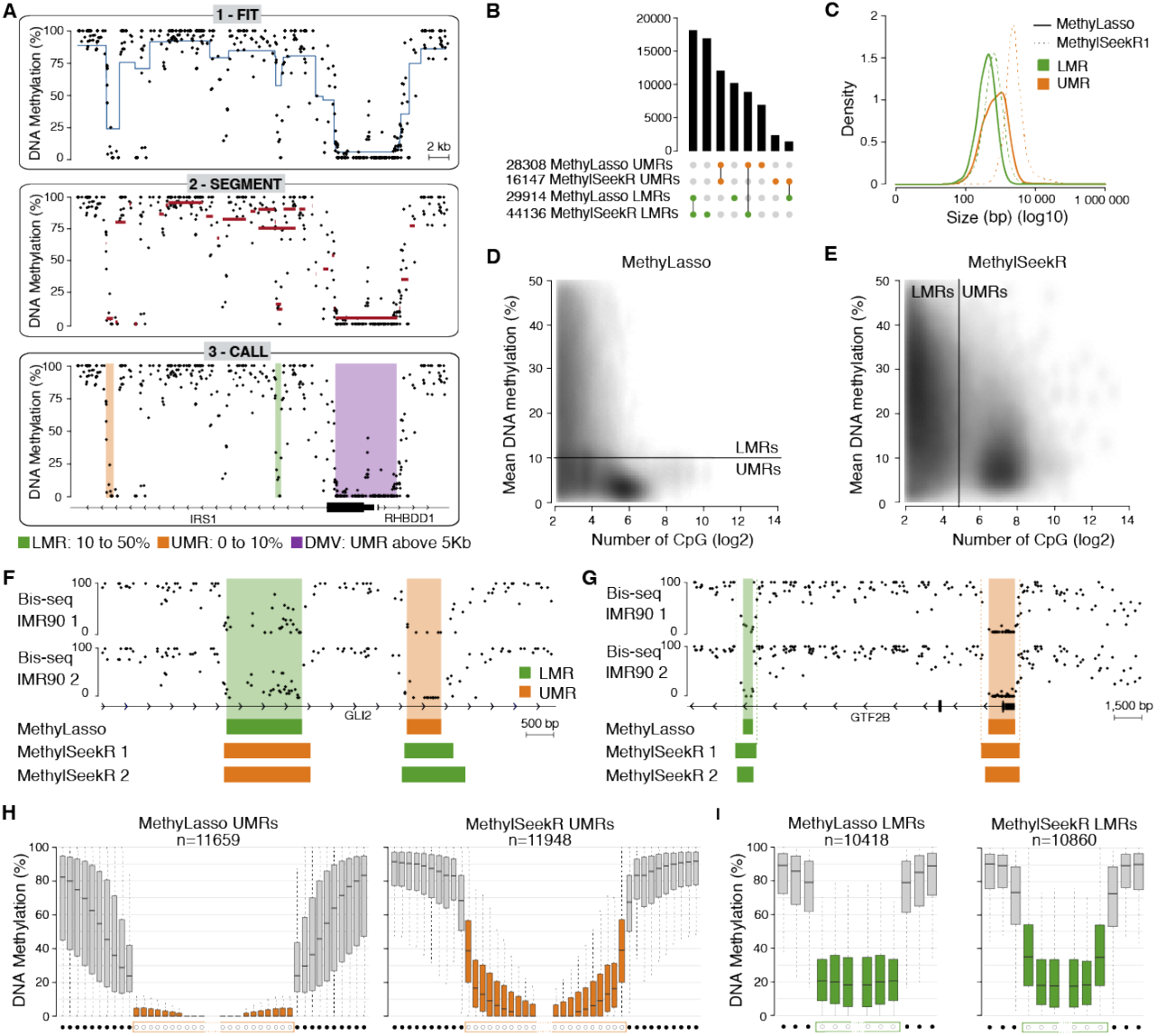
Identification of low-methylated regions (LMRs), unmethylated regions (UMRs) and DNA methylation valleys (DMVs) **A**. Genome browser view of the MethyLasso segmentation of the DNA methylation data in IMR90 cells. 1. Fit the model to the data. 2. Segment the data. 3. Call the LMRs, UMRs and DMVs. **B**. Upset plot of the number and overlap of UMRs and LMRs called by MethyLasso and MethylSeekR. MethylSeekR regions for replicate one are shown. **C**. Histogram of the size of the LMRs and UMRs called by MethyLasso and MethylSeekR. **D**. Density plot of the number of CpGs versus mean DNA methylation in each region from MethyLasso. The black line represents the MethyLasso threshold between LMRs (top) and UMRs (bottom). **E**. Density plot of the number of CpGs versus DNA methylation in each region from MethylSeekR. The black line represents the MethylSeekR threshold between LMRs (left) and UMRs (right). **F**. Genome browser view of an example of different annotations of LMRs and UMRs between MethyLasso and MethylSeekR. **G**. Genome browser view of an example of differences in boundaries between MethyLasso and MethylSeekR LMR and UMR regions. **H**. Boxplot of the DNA methylation at individual CpGs at UMR boundaries both called by MethyLasso and MethylSeekR. Grey boxes correspond to the ten CpGs upstream and downstream of the UMRs. Orange boxes correspond to the ten CpGs at the beginning or the end of the UMRs. **I**. Boxplot of the DNA methylation at individual CpGs at LMR boundaries both called by MethyLasso and MethylSeekR. Grey boxes correspond to the three CpGs upstream and downstream of the LMRs. Green boxes correspond to the three CpGs at the beginning or the end of the LMRs.

### Identification of partially methylated domains (PMDs)

MethyLasso identifies PMDs by segmenting the genome into large regions above 5 kb with methylation levels below 70% and standard deviation above 0.15 (**see Methods**). In contrast to MethylSeekR, MethyLasso does not perform a preliminary analysis defining if the data contains or not PMDs. When calling PMDs on data from human ESH1 embryonic stem cells (28) expected not to have any PMDs, MethyLasso indeed found only 61 PMDs. In human IMR90 fibroblast data, MethyLasso identified 9,740 PMDs, which are largely overlapping PMDs found by MethylSeekR (**Figure 2A,B and Supplementary Figure 3**). One advantage over MethylSeekR is that MethyLasso can integrate replicates when available to call PMDs only once for the condition. While only few MethyLasso PMDs were not identified by MethylSeekR, a large fraction of MethylSeekR PMDs were not identified by MethyLasso (**Figure 2A**). However, those have a high level of DNA methylation above 70%, which do not qualify as PMDs according to MethyLasso thresholds (**Figure 2B,C and Supplementary Figure 3**).

### Identification of differentially methylated regions (DMRs)

MethyLasso can also identify DMRs in DNA methylation data from different conditions. It fits the model to DNA methylation differences across conditions, segments the data and calls DMRs according to specific thresholds such as by default a mean methylation difference above 10% and a minimum of 4 CpGs (**see Methods**). It also annotates the DMRs using the previous segmentation from condition one to condition two e.g. LMRtoUMR, (nothing)toPMD, fromDMV(to nothing). To evaluate the performance of MethyLasso in identifying DMRs, we compared it to six other approaches: Defiant (21), DSS (20), DMRcate (23), Dmrseq (22), Radmeth (19) and Metilene (24), using Bis-seq datasets from human colon cancer compared to healthy samples (29). MethyLasso identified the most DMRs (n= 272,123) in cancer versus healthy samples (**Figure 3A**). Defiant, DSS and DMRcate also identified several hundreds of thousands of DMRs whereas Dmrseq, Radmeth and Metilene identified much fewer DMRs (only 23,141 for Metilene) (**Figure 3A**). In order to compare the different tools, we focused our analyses on the best 150,000 DMRs from each method using relaxed threshold when fewer regions were identified by default. When looking at the DMRs’ DNA methylation differences, Dmrseq and DMRcate mostly identified DMRs with small differences, MethyLasso and Defiant identified more DMRs with small differences close to the 10% threshold but also some with big differences, whereas DSS, Radmeth and Metilene identified mostly DMRs with bigger differences (**Figure 3B**). When looking at the DMR’s size, MethyLasso, Defiant and Metilene identified DMRs with a wide range of sizes whereas other approaches have a more limited range with Radmeth DMRs being short, DSS DMRs having an intermediate size and Dmrseq and DMRcate DMRs being large (**Figure 3C**). We also evaluated the DMR’s boundaries as we noticed that the DMRs identified by the different tools have varying boundaries that can extend to neighboring CpGs with lower methylation differences (**Figure 3D**). To evaluate this systematically, we compared the three first and last CpGs within DMRs to the three CpGs immediately upstream or downstream outside DMRs. We observed that MethyLasso DMRs have sharp boundaries of DNA methylation differences compared to the neighboring CpGs (**Figure 3E**) and homogenous levels of methylation differences within DMRs. Defiant DMRs also have sharp boundaries but the median methylation difference within DMRs is less homogenous. In contrast, the methylation differences at the DMR boundaries of DSS, DMRcate and Radmeth have a bell shape and the CpGs just outside their DMRs still have methylation differences higher than 10% (our threshold), indicating that their boundaries are less accurate. DMRcate and Dmrseq DMRs only have small shifts and CpGs in DMRs have lower DNA methylation differences indicating that their boundaries are not accurate. This might be due to the fact that it smooths the data before calling DMRs and identifies mainly large DMRs. For Metilene, which is also a segmentation-based approach, it’s DMRs boundaries are well defined (**Figure 3E**). Importantly, we obtained similar results when identifying DMRs in completely different datasets: human ESH1 versus IMR90 cells (28) with big DNA methylation differences, or mouse hematopoietic stem cells (HSCs) versus multipotent progenitors (MPPs) (30) that show small changes in DNA methylation (**Supplementary Figure 4**). Altogether, these analyses show that MethyLasso identifies a higher number of DMRs with clear boundaries compared to existing DMR tools.

**Figure 2.**
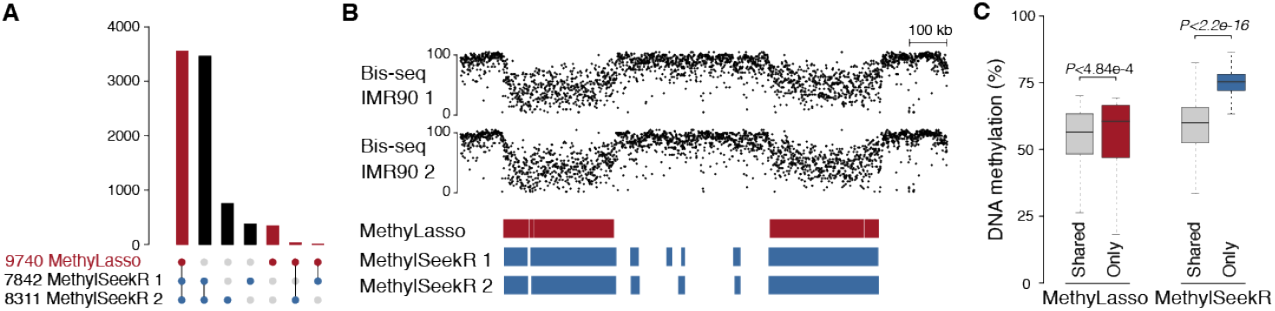
Identification of partially methylated domains (PMDs) **A**. Upset plot of the number and overlap of PMDs from IMR90 cells called by MethyLasso and MethylSeekR. MethylSeekR PMDs called independently for the two replicates are shown. **B**. Genome browser view of an example of PMDs shared by both MethyLasso and MethylSeekR or only found by MethylSeekR. Region from hg38 at chr3:154,200,000-155,900,000. **C**. Boxplot of the mean DNA methylation of the MethyLasso PMDs shared with MethylSeekR (grey) or not (red) and of the MethylSeekR PMDs shared with MethyLasso (grey) or not (blue). Corresponding Wilcoxon p-value.

**Figure 3.**
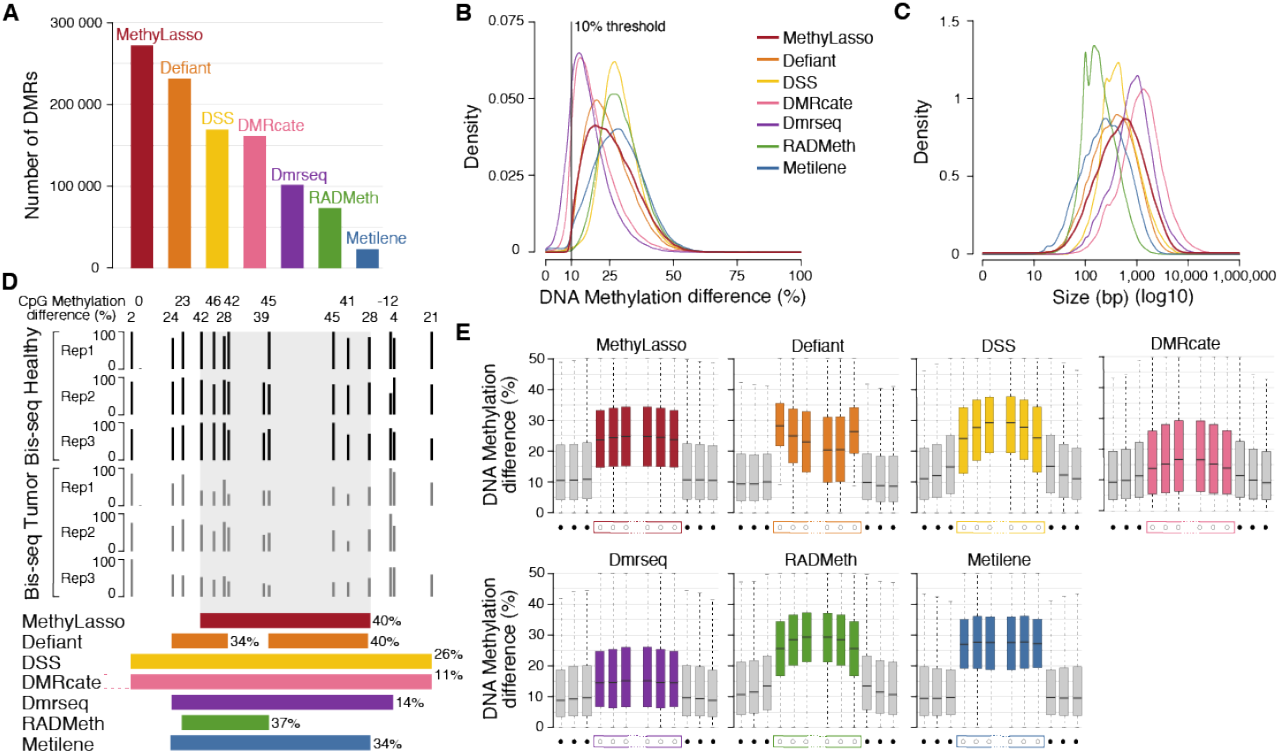
Identification of differentially methylated regions (DMRs) **A**. Barplot of the number of DMRs between healthy and colon cancer cells identified by the different methods. **B**. Histogram of the absolute DNA methylation difference in the best 150,000 DMRs called by the different methods. **C**. Histogram of the size of the best 150,000 DMRs called by the different methods. **D**. Genome browser view of an example of DMRs with different region boundaries found by all methods. DMR from DMRcate extend much upstream of the region shown (chr5:72417868-72419158). Methylation difference is shown in percent for all individual CpGs and for DMRs from each method. Region from hg38 at chr5:72,418,800-72,419,200. **E**. Boxplot of the difference of DNA methylation at individual CpGs at the best 150,000 DMR region boundaries called by the different methods. Grey boxes correspond to the three CpGs upstream and downstream of the DMRs. Colored boxes correspond to the three CpGs at the beginning or the end of the DMRs. Comparisons between CpGs located out/grey vs. in/red DMRs are significant for all approaches Wilcoxon p-value < 2.2e-16

### Consistency of DMRs from different approaches

Next, we investigated the consistency of MethyLasso DMRs compared to the other methods. To compare the DMRs from the different approaches, we first analyzed the overlap between the best 150,000 DMRs identified by any approach merged into 386,580 regions. Most are shared by at least two approaches (69%) but only 5% are shared by all and a substantial fraction is found by only one method (31%) (**Figure 4A,B**). As expected, the fraction of DMRs shared by all and therefore likely to be true increased with an increasing methylation difference, with a strong improvement starting at 20% methylation difference (**Supplementary Figure 5**). However, the evaluation of the DMR’s overlap might be underestimated since some might be shared but with ranks slightly below our threshold (34). Therefore, we asked whether the very best 50,000 DMRs from one approach were identified by the best 150,000 DMRs of the other approaches. Out of the best 50,000 DMRs identified by MethyLasso, 18.5% were found by all other approaches but 81% were found by some others and only 0.5% were not found by any other approach (**Figure 4C** red, **Supplementary Figure 6** for examples and all data available for visualization in a UCSC hub, see methods). When looking at the difference in methylation of those DMRs, we observed that the DMRs found by MethyLasso and others had bigger methylation differences than the ones found by Methylasso alone (**Figure 4D** red). This might be explained by the fact that DMRs with small changes in DNA methylation are more difficult to identify. Nevertheless, DMRs found by MethyLasso alone had a similar CpG content than the DMRs found by MethyLasso and others (including some with many CpGs) (**Figure 4E** red). This suggests that MethyLasso DMRs are likely true. When looking at the best 50,000 DMRs identified by others compared the whole 150,000 sets, most were also identified by MethyLasso (**Figure 4C**; 95.1% for Metilene, 87.4% for DSS, 66.7% for RADmeth, 65.5 for DMRcate, 62.7% for Dmrseq and 60.1% for Defiant). We then analyzed the DMRs found by these other methods but not MethyLasso. Defiant DMRs not found by MethyLasso had low methylation differences (**Figure 4D** orange) and were enriched in DMRs with a small number of CpGs (between 4 and 6, **Figure 4E** orange), which could be artifacts. DSS DMRs not found by MethyLasso do not have smaller methylation differences (**Figure 4D** yellow) and lower CpG numbers (**Figure 4E** yellow), but we observed that they had a low coverage of their CpGs below the 70% coverage threshold set by MethyLasso (**Figure 4F** yellow and **Supplementary Figure 6** for examples and all data available for visualization in a UCSC hub, see methods). Most DMRs from DMRcate are large regions (**Figure 3C**, pink) with low methylation differences (**Figure 4D**, pink) but some of the ones not identified by MethyLasso have a low number of CpGs (**Figure 4E** pink) and a low coverage of their CpGs (**Figure 4F** pink). Dmrseq DMRs are also large regions (**Figure 3C**, purple) and have overall low methylation differences that are even lower when not found by MethyLasso (**Figure 4D** purple) but still have similar number of CpGs (**Figure 4E** purple). Radmeth DMRs that are small in size (**Figure 3C**, green) have overall higher methylation difference but DMRs not found by any others have lower methylation differences (**Figure 4D** green). However, most of Radmeth DMRs not found by MethyLasso have a very low number of CpGs (**Figure 4E** green), which could be artifacts. Finally, only very few Metilene DMRs were not found by MethyLasso (**Figure 4E** blue). Some have a low coverage of its CpGs (**Figure 4F** blue) and 59 have no coverage at all for one of the conditions (**Supplementary Figure 6** for examples), which is due to the fact that Metilene imputes missing data. Importantly, we obtained similar results when identifying DMRs in completely different cell lines namely human ESH1 versus IMR90 cells (28) with bigger DNA methylation changes or in differentiating cells namely mouse HSC versus MPP cells (30) with small changes in DNA methylation (**Supplementary Figures 7 & 8**). Altogether, these analyses show that MethyLasso identifies DMRs that are also identified by others, and that DMRs from others not identified by MethyLasso tend to have features that could come from artifacts.

**Figure 4.**
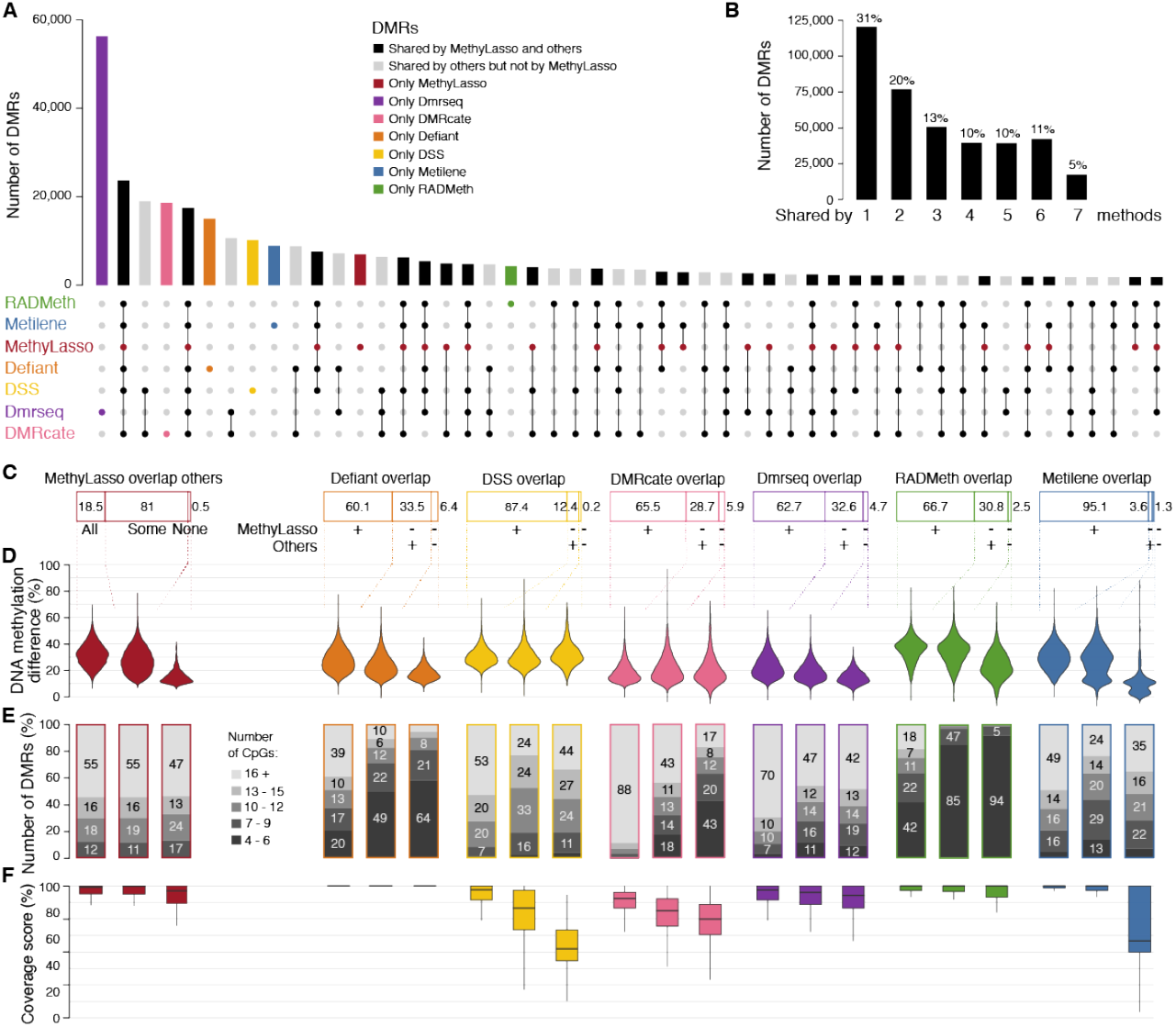
Consistency between DMRs from different approaches. **A**. Upset plot of the overlap of best 150,000 DMRs called by the different methods merged into 386,580 regions. Bars corresponding to DMRs identified by MethyLasso are represented in red. **B**. Barplot summarizing the upset plot by summing the number of regions identified by all, some or none of the methods. **C**. Cumulative barplot showing the percent of best 50,000 MethyLasso DMRs overlapping with all others 150,000 DMRs, some others or none of the other methods. For other methods, percent of their best 50,000 DMRs overlapping with 150,000 DMRs from MethyLasso (+), not MethyLasso but others (-/+) or not MethyLasso nor others (-/-). **D**. Absolute DNA methylation difference in the DMRs from the categories in C. **E**. Cumulative barplot showing the number of CpGs in the DMRs from the categories in C. **F**. Boxplot showing the coverage of CpGs in the DMRs from the categories in C.

### Sensitivity and precision of the approaches using simulated DMRs

To evaluate the performance of MethyLasso at identifying DMRs and to compare it to other approaches, we called DMRs on simulated data with varying levels of DNA methylation differences (data from Metilene, **see Methods**). We first investigated the sensitivity of the approach, which evaluates the fraction of simulated DMRs that are correctly identified or true positive rate (**Figure 5A**). For DMRs with large methylation differences (40 to 60%), which should be the easiest to identify, only MethyLasso and Metilene were able to identify all simulated DMRs (sensitivity at 1). For DMRs with small methylation differences (10 to 20%), which should be more difficult to identify, Defiant and MethyLasso performed best followed by Metilene, whereas Dmrseq, DSS, Radmeth and DMRcate performed worse. We then investigated the precision of the approach, which evaluates the fraction of predicted DMRs that were indeed simulated (**Figure 5B**). For DMRs with large methylation differences, all programs identified well the simulated DMRs. For DMRs with small methylation differences, all programs performed well with Radmeth being best closely followed by DSS, DMRcate, Metilene, Dmrseq and MethyLasso and Defiant being less precise. Finally, we calculated the F1-score i.e. the harmonic mean of the sensitivity and precision and identified MethyLasso and Metilene as performing perfectly on DMRs with large differences and MethyLasso followed by Metilene and then Defiant on DMRs with small differences (**Figure 5C**). Performances and ranking are similar when a different set of simulated DMRs are used (**Supplementary Figure 9**). In summary, we conclude that MethyLasso outperforms existing tools on the analysis of simulated DMRs.

**Figure 5.**
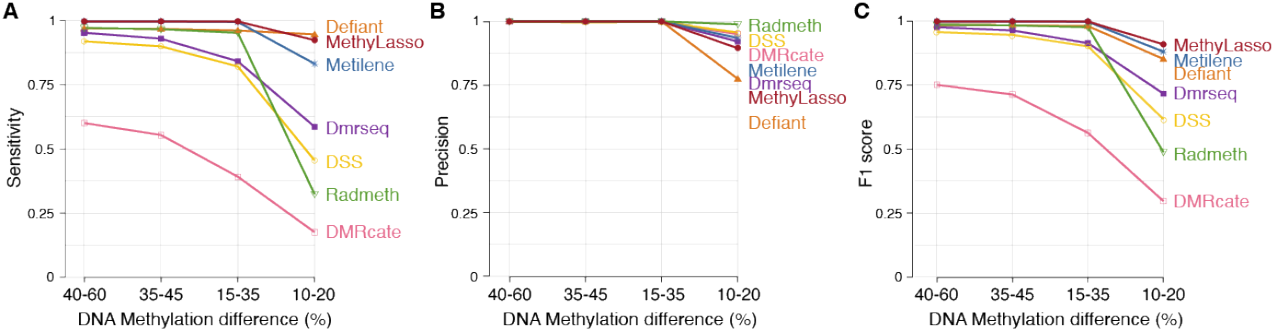
Sensitivity and precision of the method using simulated DMRs. Simulated DMRs in different bins of DNA methylation difference from Metilene using a heterogeneous background. **A**. Sensitivity or recall of the predictions measured as the number of true positives among all simulated. **B**. Precision of the predictions measured as the number of true positives among all predicted. **C**. F1 score or accuracy of the predictions measured using both the sensitivity and precision.

## DISCUSSION

MethyLasso is a new approach to analyze DNA methylation patterns from whole-genome Bis-seq datasets. It mostly differs from other tools as it is based on a segmentation of the DNA methylation patterns. MethyLasso can analyze data from each condition independently and can integrate replicates to identify LMRs, UMRs and DMVs by performing a segmentation of the DNA methylation levels, and PMDs by performing a segmentation of the DNA methylation variation. It differs from MethylSeekR mainly by defining regions of interest based on their DNA methylation levels matching their definition (LMR or UMR) rather than based on their CpG content (**Figure 1D,E**). Additionally, we show that MethyLasso performs better at defining the region’s boundaries since it does not smooth methylation levels (**Figure 1H,I**). To identify PMDs, MethyLasso focuses on large regions with heterogeneous DNA methylation levels and does not identify MethylSeekR PMDs with high and stable methylation levels (**Figure 2B,C**).

MethyLasso can also be applied to compare DNA methylation levels across two conditions and can integrate replicates to identify DMRs by performing a segmentation of the DNA methylation differences that defines DMRs based on the overall methylation patterns rather than grouping individual CpGs. When comparing different tools, the DMRs identified have a limited overlap (**Figure 4A,B**) indicating that the identification of DMRs is a challenging task. As expected, DMRs with bigger DNA methylation changes have a higher overlap than DMRs with small changes (**Supplementary Figure 5**).

MethyLasso identifies a large number of DMRs, some with big DNA methylation changes and more as expected with small changes close to the cutoff (**Figure 3A,B**). MethyLasso DMRs have a wide range of sizes showing that it can adapt well to the size of potential DMRs (**Figure 3C**). The boundaries of the MethyLasso DMRs are well-defined at locations with sharp transitions of DNA methylation changes with all CpGs within DMRs having stable high levels of DNA methylation and CpGs just outside having stable levels close to the 10% cutoff (**Figure 3E**). 99% of the best MethyLasso DMRs are identified by at least one other method indicating that they are most likely true positives (**Figure 4C**). The few DMRs not identified by any of the others have small methylation changes and are therefore more difficult to identify. Between 95% (Metilene) and 60% (Defiant) of the best DMRs from other methods are identified by MethyLasso. Most of the ones missed by MethyLasso have small methylation changes. DMRs identified by Defiant, RADmeth and DMRcate and not by MethyLasso are mainly DMRs containing few CpGs (4 to 6). Some DMRs identified by DSS, DMRcate and Metilene but missed by MethyLasso still have high levels of methylation changes but come from regions that are not well covered (DSS) or where data was imputed (Metilene). Finally, using simulated DMRs with different bins of DNA methylation change, MethyLasso is the approach with the best overall results especially in term of sensitivity where it identifies best the simulated DMRs with both high and low methylation changes (**Figure 5**).

Defiant DMRs have a wide range of DNA methylation differences and size and good region boundaries. However, they have the least overlap with MethyLasso ones (60%), the most not found by any others (6.4%) and those have only few CpGs (**Figure 4C,E**), all of which indicates that they could be artifacts. On simulated data, its sensitivity is high but it surprisingly misses DMRs with big methylation changes and its precision is lower than all others (**Figure 5**).

DSS tends to identify DMRs with bigger methylation changes but less well-defined region boundaries that might be due to smoothing (**Figure 3B, E**). It also includes regions with low CpG coverage that might bias the methylation levels and represent false positives (**Figure 4F** and **Supplementary Figure 6**). On simulated data, DSS has a poor sensitivity (**Figure 5**).

DMRcate identifies large DMRs and does not adapt well to short regions of DNA methylation changes resulting in DMRs with small methylation changes (**Figure 3B,C**). It’s DMR boundaries less well-defined, which might be due to smoothing (**Figure 3E**). On simulated data, DMRcate shows to have the worse sensitivity (**Figure 5**).

Dmrseq also identifies large DMRs with small methylation changes and not well-defined boundaries, which might be due to smoothing (**Figure 3B,C,E**). On simulated data, Dmrseq shows to have a poor sensitivity (**Figure 5**). Additionally, Dmrseq can only be applied if replicates are available.

RADmeth identifies a low number of DMRs with small size and therefore less well-defined region boundaries (**Figure 3B,C,E**). On simulated data, RADmeth shows to have a poor sensitivity (**Figure 5**).

Metilene only identifies very few DMRs (more than 10 times less than MethyLasso) with big methylation changes even though the threshold is set to 10% (**Figure 3A,B**). Like MethyLasso, Metilene applies a segmentation approach resulting in well-defined region boundaries. The overlap between Metilene and MethyLasso DMRs is very high and the few DMRs only identified by Metilene are mostly due to the fact that it imputes missing data, generating DMRs with artificially high methylation changes (**Figure 4F**). On simulated data that were generated by the authors of Metilene where it might have an advantage, Metilene performs worse than MethyLasso in terms of sensitivity and a slightly better in terms of precision (**Figure 5**).

In conclusion, MethyLasso applies a robust segmentation approach to analyze DNA methylation patterns either in a single condition to identify LMRs, UMRs, DMVs and PMDs or by comparing conditions to identify DMRs. We conducted an extensive benchmark to show that MethyLasso performs best compared to state-of-the-art tools. Since DNA methylation levels anticorrelate with chromatin accessibility, the identification of LMRs, UMRs and DMRs is a powerful approach to predict active regulatory regions bound by transcription factors that regulate gene expression.

## CODE AVAILABILITY

MethyLasso is available on github from https://github.com/abardet/methylasso

## ACKOWLEDGEMENTS

The authors thank Michaël Dumas and Yahia Hadj-Arab for testing MethyLasso, Nacho Molina and Pierre-Éric Lutz for feedback on the analyses and manuscript and the Weber and Molina labs for helpful discussions.

## AUTHOR CONTRIBUTIONS

Y.G.S., A.F.B., D.B. and M.W. designed the project. Y.G.S. developed the statistical model and heuristics of the method. D.B. developed the heuristics and benchmarked the method. A.F.B. and M.W. supervised the project. A.F.B., D.B., Y.G.S. and M.W. wrote the manuscript. All authors read and approved the final manuscript.

