## Supplementary Figures for "MethyLasso: a segmentation approach to analyze DNA methylation patterns and identify differentially methylation regions from whole-genome datasets"

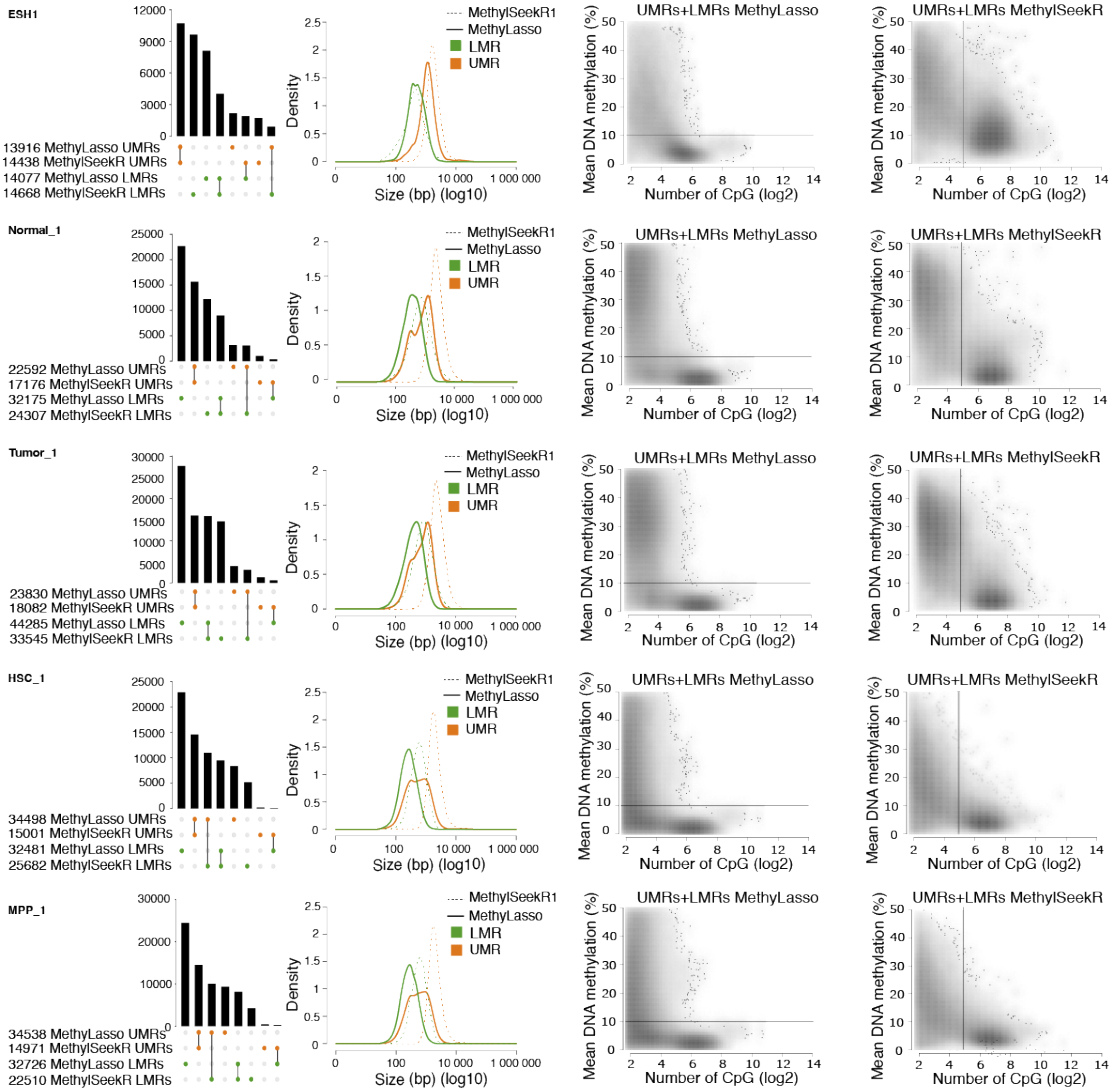

***Supplementary Figure 1 - Features of unmethylated regions (UMRs) and low-methylated regions (LMRs)***

*Same as Figure 1B, C, D, E but for different datasets.* ***A.*** *Upset plot of the number and overlap of UMRs and LMRs called by MethyLasso and MethylSeekR. MethylSeekR regions for replicate one are shown.* ***B.*** *Histogram of the size of the LMRs and UMRs called by MethyLasso and MethylSeekR.* ***C.*** *Density plot of the number of CpGs versus mean DNA methylation in each region from MethyLasso. The black line represents the MethyLasso threshold between LMRs (top) and UMRs (bottom).* ***D.*** *Density plot of the number of CpGs versus DNA methylation in each region from MethylSeekR. The black line represents the MethylSeekR threshold between LMRs (left) and UMRs (right).*

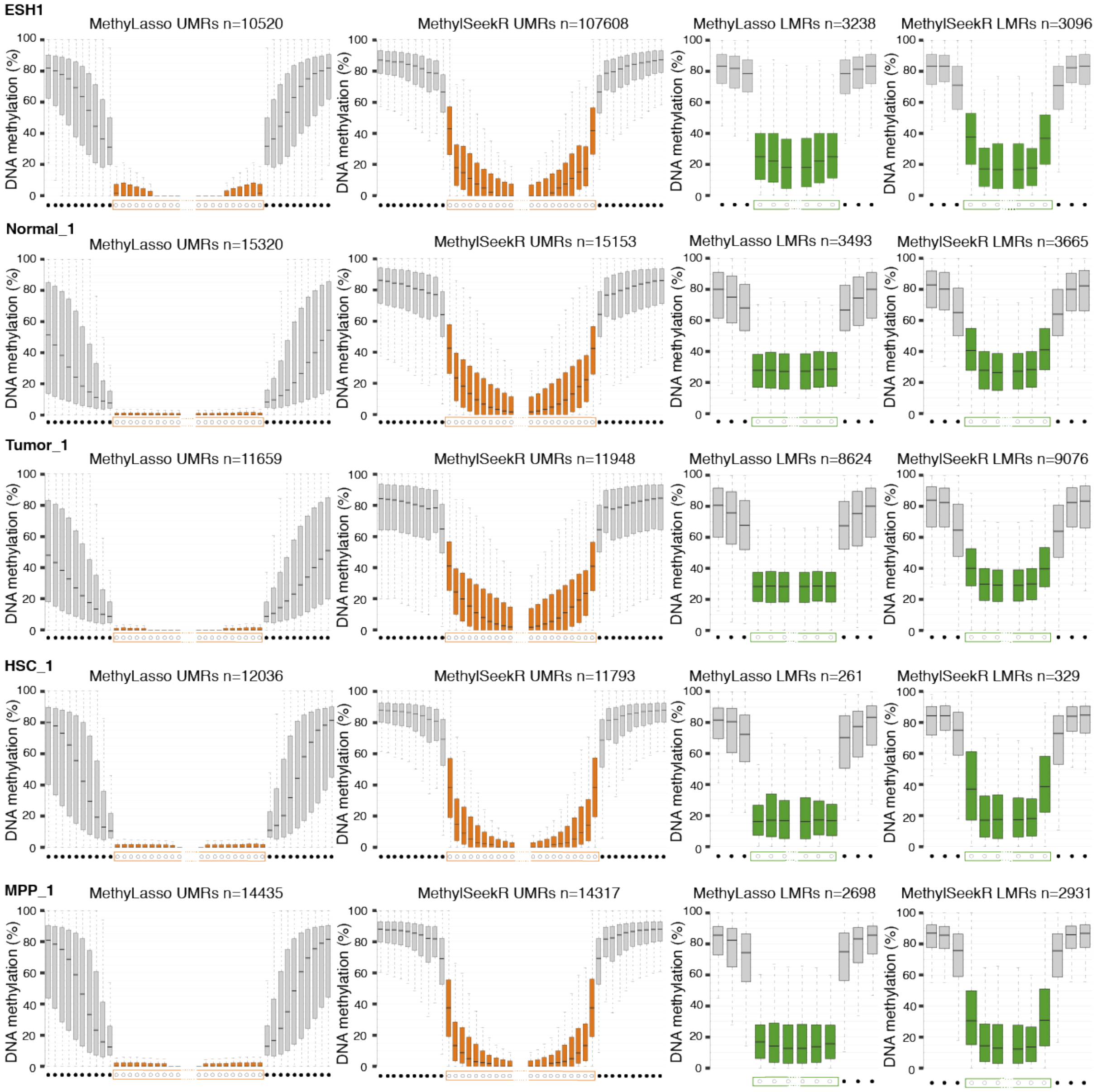


***Supplementary Figure 2 - Boundaries of unmethylated regions (UMRs) and low-methylated regions (LMRs)***

*Same as Figure 1H and I but for different datasets.* ***A.*** *Boxplot of the DNA methylation levels at individual CpGs at UMR boundaries called by MethyLasso and MethylSeekR. Grey boxes correspond to the ten CpGs upstream and downstream of the UMRs. Orange boxes correspond to the ten CpGs at the beginning or the end of the UMRs.* ***B.*** *Boxplot of the DNA methylation at individual CpGs at LMR boundaries called by MethyLasso and MethylSeekR. Grey boxes correspond to the three CpGs upstream and downstream of the LMRs. Green boxes correspond to the three CpGs at the beginning or the end of the LMRs.*

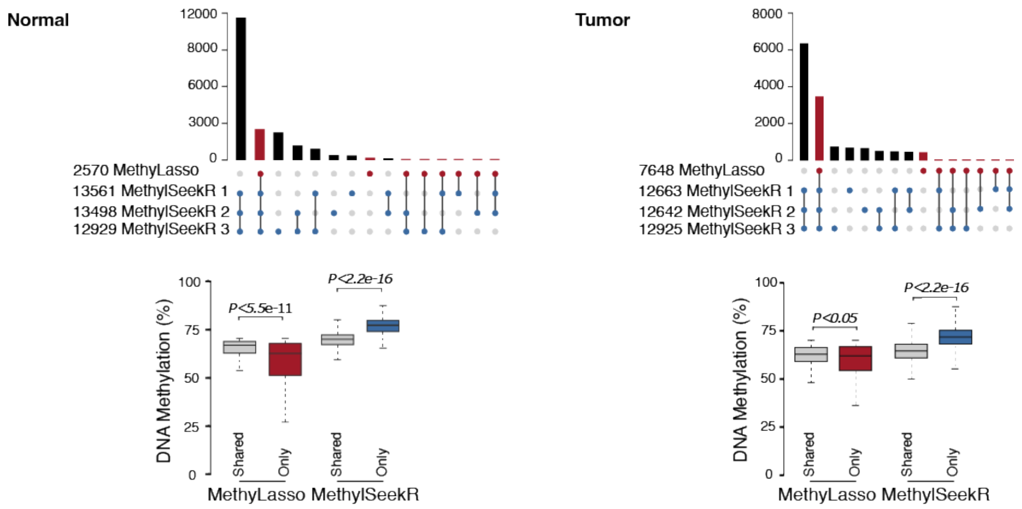


***Supplementary Figure 3 - Features of partially methylated domains (PMDs)***

*Same as Figure 2A and C but for different datasets.* ***A.*** *Upset plot of the number and overlap of PMDs called by MethyLasso and MethylSeekR. MethylSeekR PMDs called independently for the two replicates are shown.* ***B.*** *Boxplot of the mean DNA methylation of the MethyLasso PMDs shared with MethylSeekR (grey) or not (red) and of the MethylSeekR PMDs shared with MethyLasso (grey) or not (blue). Corresponding Wilcoxon p-value.*

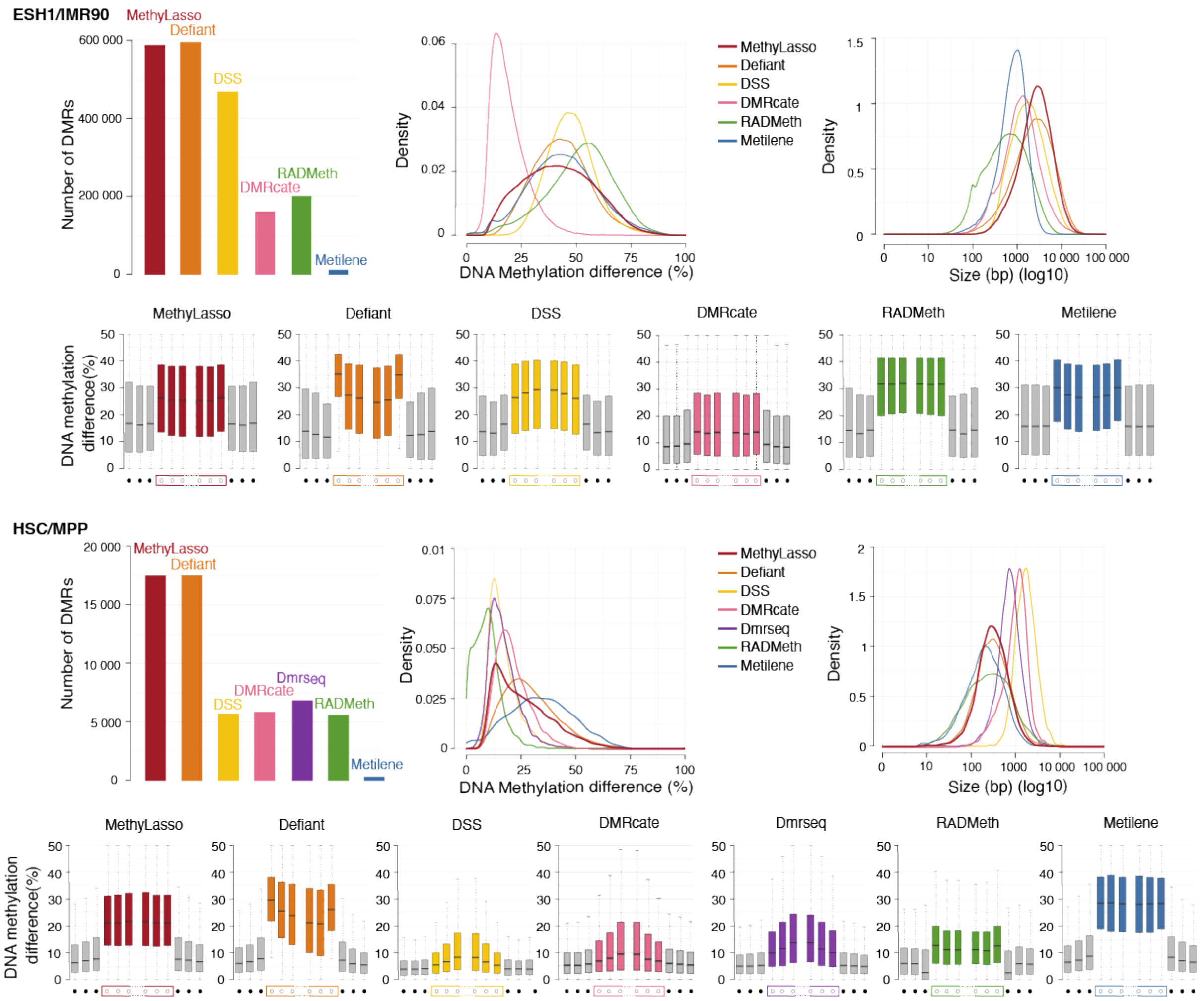


***Supplementary Figure 4 - Features of differentially methylated regions (DMRs)***

*Same as Figure 3 but for the best 270,000 DMRs for ESH1 vs. IMR90 (**A-D**) and the best 5,000 DMRs for HSC vs. MPP (**E-H**). **A&E.** Barplot of the number of DMRs identified by the different methods. **B&F.** Histogram of the absolute DNA methylation difference in the best DMRs called by the different methods. **C&G.** Histogram of the size of the best called by the different methods. **D&H.** Boxplot of the difference of DNA methylation at individual CpGs at DMR region boundaries called by the different methods. Grey boxes correspond to the three CpGs upstream and downstream of the DMRs. Colored boxes correspond to the three CpGs at the beginning or the end of the DMRs. Note that DMRs could not be identified by Dmrseq for ESH1 vs IMR90 data since it had no replicate for the ESH1 condition.*

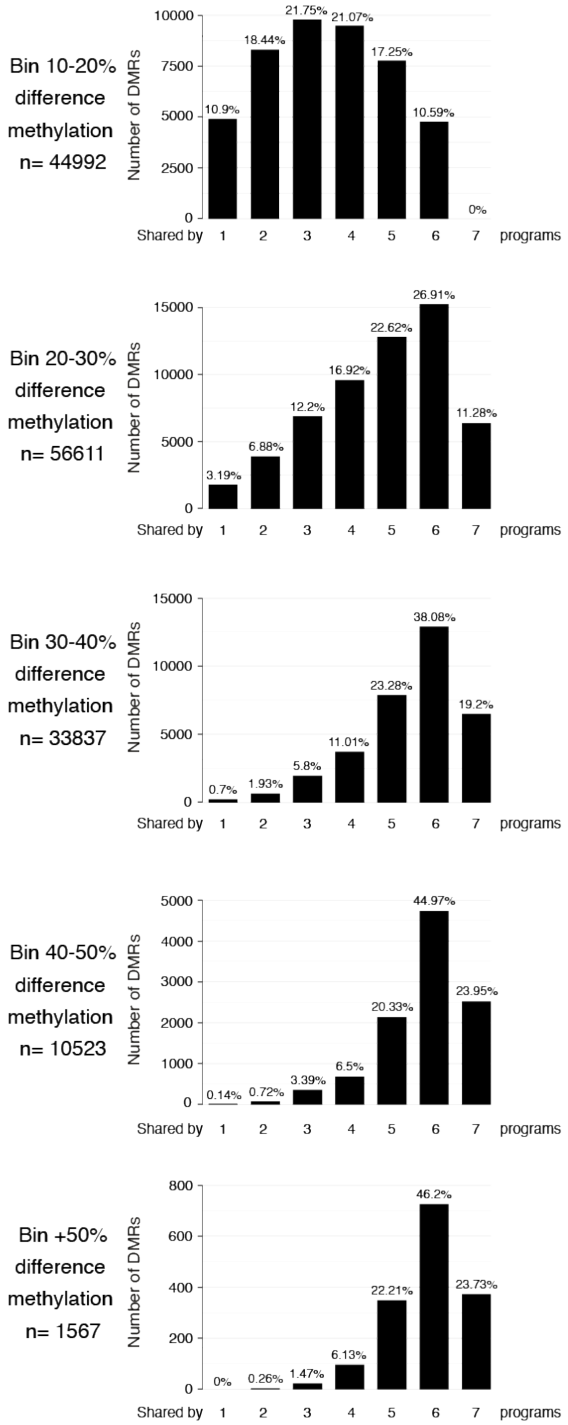


***Supplementary Figure 5 - Consistency between DMRs from different approaches at different methylation difference levels***

*Barplots summing the number of regions identified by all, some or none of the methods for different bins of methylation difference. Data from human colon cancer compared to healthy samples.*

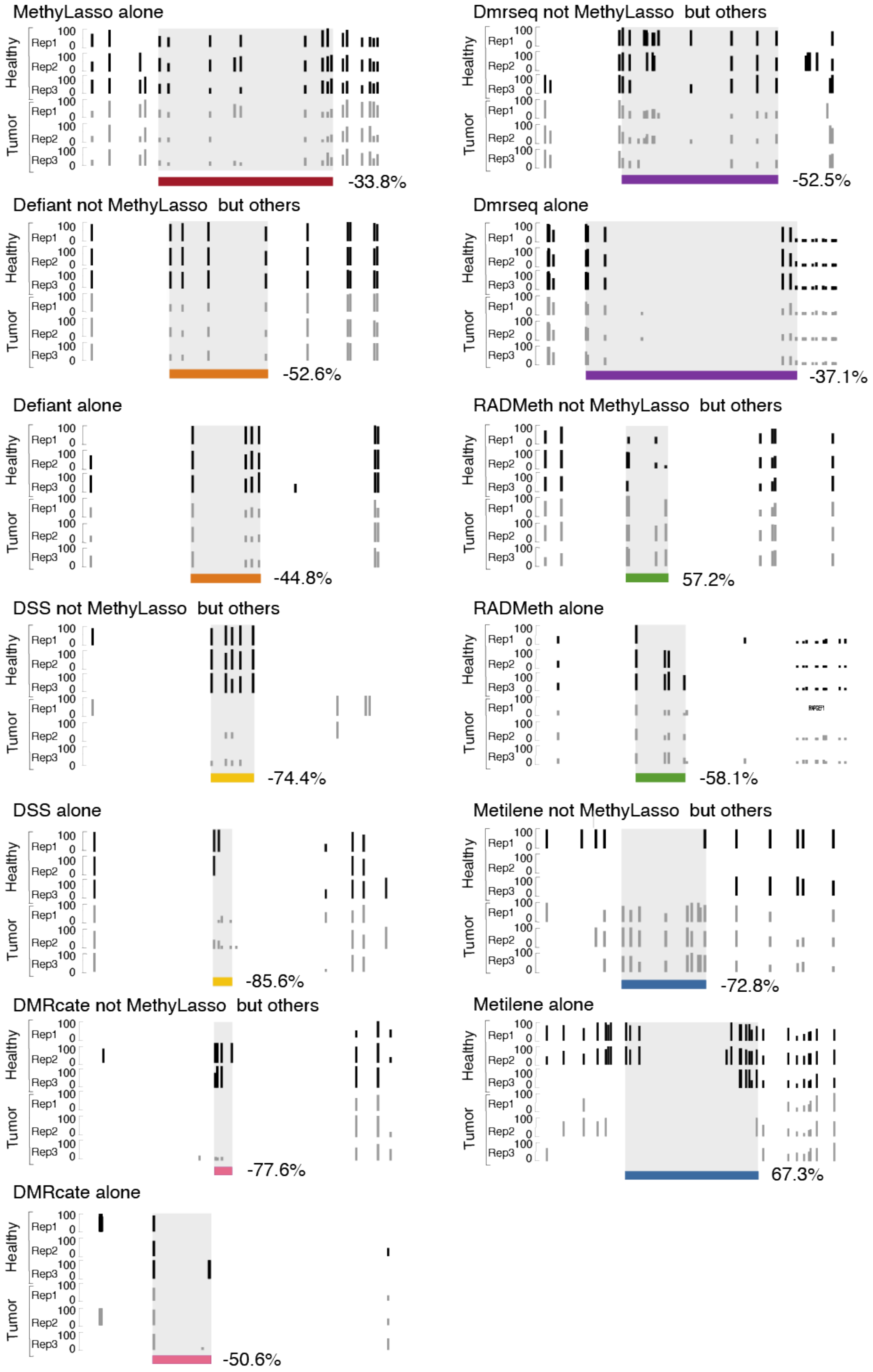


***Supplementary Figure 6 - Example of DMRs from the different methods***

*Examples of DMRs from each category in Figure 4C chosen among the top ten.*

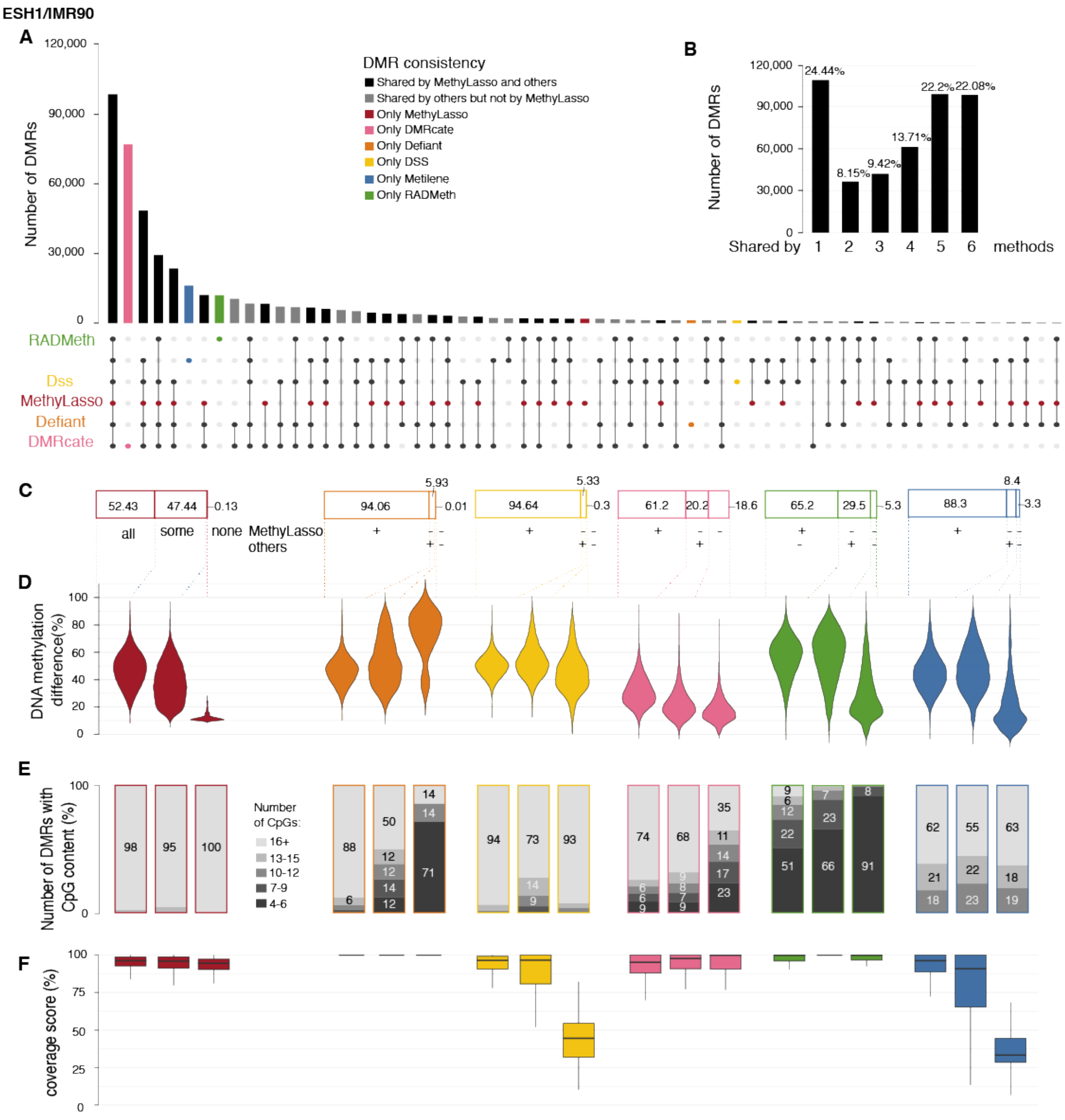


***Supplementary Figure 7 - Consistency between DMRs from different approaches in ESH1 vs. IMR90 data***

*Same as Figure 4.* ***A.*** *Upset plot of the overlap of best 270,000 DMRs called by the different methods. Bars corresponding to DMRs identified by MethyLasso are represented in red.* ***B.*** *Barplot summarizing the upset plot by summing the number of regions identified by all, some or none of the methods.* ***C.*** *Cumulative barplot showing the percent of best 100,000 MethyLasso DMRs overlapping with all others 270,000 DMRs, some others or none of the other methods. For other methods, percent of their best 100,000 DMRs overlapping with 270,000 DMRs from MethyLasso (+), not MethyLasso but others (-/+) or not MethyLasso nor others (-/-).* ***D.*** *Absolute DNA methylation difference in the DMRs from the categories in C.* ***E.*** *Cumulative barplot showing the number of CpGs in the DMRs from the categories in C.* ***F.*** *Boxplot showing the coverage of CpGs in the DMRs from the categories in C.*

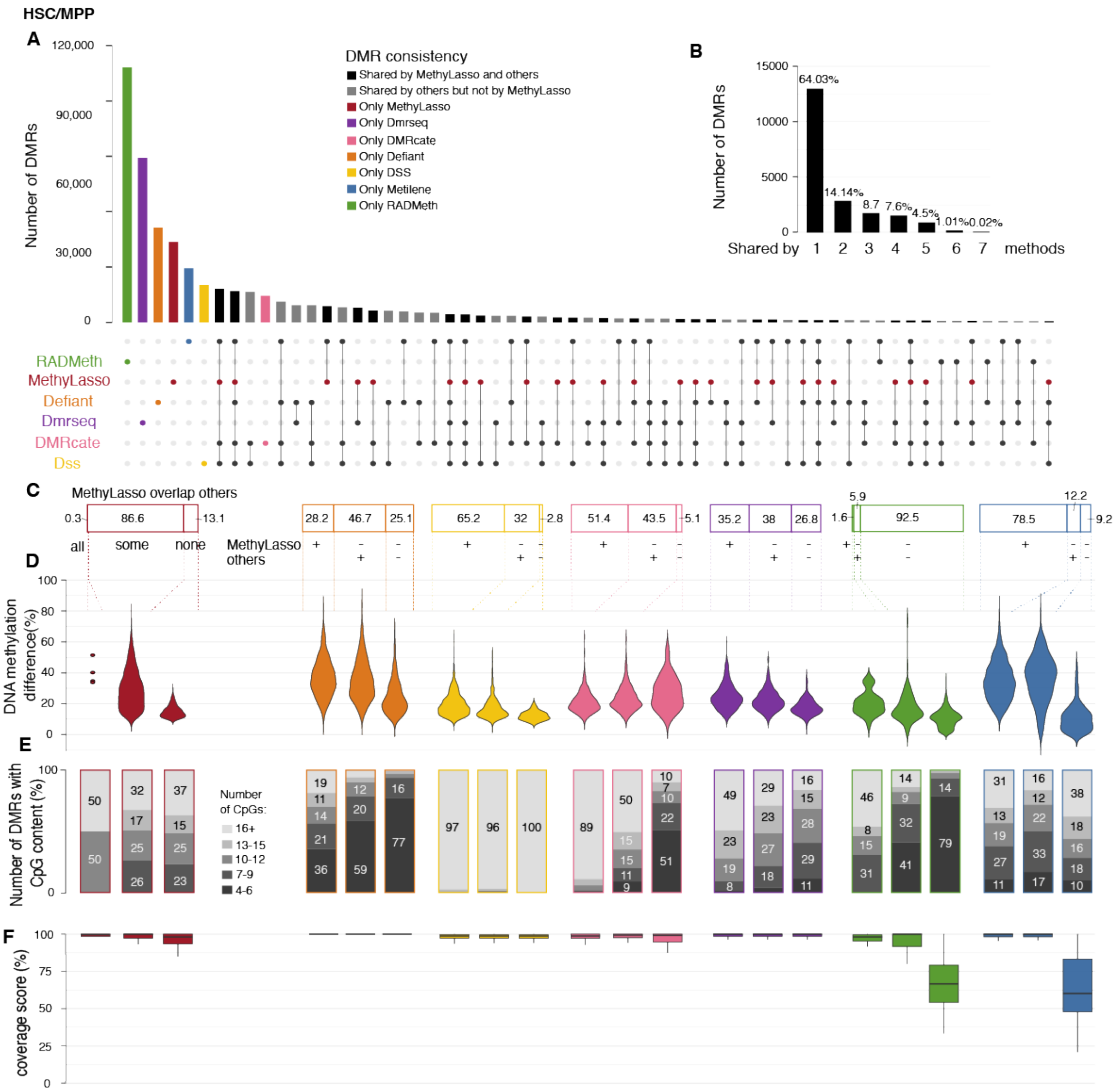


***Supplementary Figure 8 - Consistency between DMRs from different approaches in HSC vs. MPP data***
*Same as Figure 4. **A.** Upset plot of the overlap of best 5,000 DMRs called by the different methods. Bars corresponding to DMRs identified by MethyLasso are represented in red. **B.** Barplot summarizing the upset plot by summing the number of regions identified by all, some or none of the methods. **C.** Cumulative barplot showing the percent of best 1,600 MethyLasso DMRs overlapping with all others 5,000 DMRs, some others or none of the other methods. For other methods, percent of their best 1,600 DMRs overlapping with 5,000 DMRs from MethyLasso (+), not MethyLasso but others (-/+) or not MethyLasso nor others (-/-). **D.** Absolute DNA methylation difference in the DMRs from the categories in C. **E.** Cumulative barplot showing the number of CpGs in the DMRs from the categories in C. **F.** Boxplot showing the coverage of CpGs in the DMRs from the categories in C.*

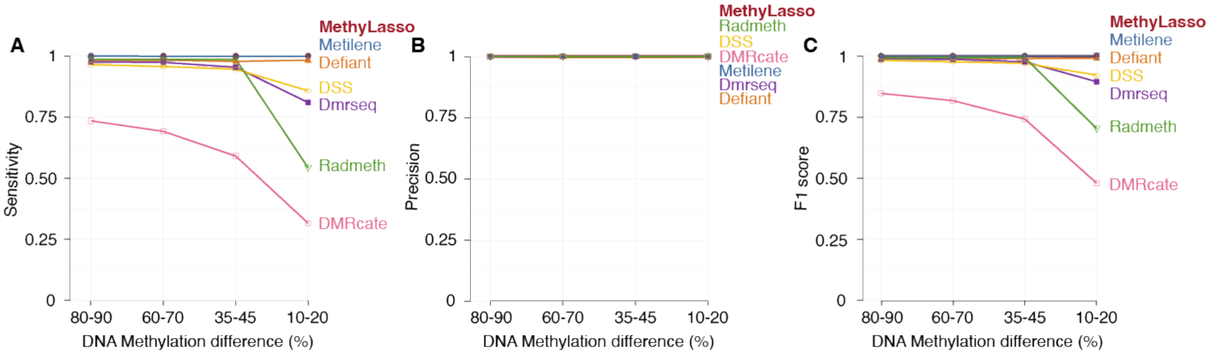


***Supplementary Figure 9 - Sensitivity and precision of the method using simulated DMRs***

*Simulated DMRs in different bins of DNA methylation difference from Metilene using a homogeneous background.* ***A.*** *Sensitivity or recall of the predictions measured as the number of true positives among all simulated.* ***B.*** *Precision of the predictions measured as the number of true positives among all predicted.* ***C.*** *F1 score or accuracy of the predictions measured using both the sensitivity and precision.*
